# Shared Hierarchical Representations Explain Temporal Correspondence Between Brain Activity and Deep Neural Networks

**DOI:** 10.1101/2025.05.19.655003

**Authors:** Eric Lützow Holm, Giovanni Marraffini, Diego Fernandez Slezak, Enzo Tagliazucchi

## Abstract

The visual cortex and artificial neural networks both process images hierarchically, progressing from low-level features to high-level semantic representations. We investigated the temporal correspondence between activations from multiple convolutional and transformer-based neural network models (AlexNet, MoCo, ResNet-50, VGG-19, and ViT) and human EEG responses recorded during visual perception tasks. Leveraging two EEG datasets of images presented at different durations, we assessed whether this correspondence reflects general architectural principles or model-specific computations. Our analysis revealed a robust mapping: early EEG components correlated with activations from initial network layers and low-level visual features, whereas later components aligned with deeper layers and semantic content. Moreover, for images presented during longer times, the extent of correspondence correlated with the semantic contribution to the EEG response. These findings highlight a consistent temporal alignment between biological and artificial vision, suggesting that this correspondence is primarily driven by the hierarchical transformation of visual to semantic representations rather than by idiosyncratic computational features of individual network architectures.

## 1 Introduction

Humans interact with a wide variety of objects in daily life, recognizing and generalizing complex sensory patterns. Object recognition relies on hierarchical neural processing in the ventral visual pathway, where neurons progressively transform low-level visual input into abstract semantic representations [1, 2]. The neural dynamics underlying this transformation have been studied using non-invasive techniques such as electroencephalography (EEG) and magnetoencephalography (MEG) [3]. Early evoked neural activity primarily reflects low-level image features—such as luminance, contrast, orientation, and color—while later components encode object identity and conceptual content, indicating a neural semantic representation [4].

Artificial neural networks (ANNs) that achieve high performance on object recognition tasks exhibit a similar hierarchical organization, with multiple layers transforming visual features at increasing levels of abstraction [5]. While the specific computations differ across models, human neuroimaging functional magnetic resonance imaging (fMRI) studies have revealed anatomical correspondences between biological and artificial representations during object recognition [6, 7, 8, 9, 10, 11]. Similar results have been obtained in monkey electrophysiological studies [12, 9]. These correspondences also extend to the temporal domain: early brain responses (measured via EEG/MEG) are best predicted by early ANN layers, while late responses align with deeper layers [13, 14, 15].

The representational similarity between brain and ANN representations depends on multiple factors, including training data, classification task and performance [6]. Importantly, this similarity does not imply that artificial systems mimic the brain’s computational mechanisms. Human fMRI and macaque electrophysiology studies show that widely different ANN models are in correspondence with brain activity along the ventral pathway, regardless of their biological interpretability [16]. For example, convolutional neural networks (CNNs) share several similarities with cortical processing, with a hierarchy of neurons presenting increasingly large receptive fields [17, 2]. Other models, such as Vision Transformers (ViTs) and multilayer perceptrons (MLPs), employ computations with a less clear neurobiological interpretation. However, both have been equally supported as plausible models to predict fMRI activity during object perception and recognition [16, 6].

We tested the hypothesis that brain-ANN temporal correspondence reflects two aspects commonly found in artificial vision models: the hierarchical organization of DNNs [18] and their transformation of low-level visual features to semantic representations [19]. Using representational similarity analysis (RSA) [20], we compared EEG-based decoding similarities with layer-specific representational similarities extracted from five object-recognition models (AlexNet [21], MoCo [22], ResNet-50 [23], VGG-19 [24], and ViT [25]). Leveraging two independent EEG datasets allowed us to investigate the effect of stimulus duration (50 ms vs. 150 ms) on brain-ANN temporal correspondence. Finally, we investigated how this correspondence depended on the contribution of visual and semantic content to ANN activations and EEG responses.

## 2 Related Work

### Brain-ANN Temporal Correspondence

Extensive fMRI work revealed a hierarchical correspondence between multiple ANNs and anatomical areas involved in vision and object recognition [8, 9, 10, 11, 6, 16]. CNNs capture the dynamics of human visual processing along the dorsal and ventral streams, as revealed using MEG [13, 15]. Deeper layers of the CNN predict late EEG responses and vice-versa for early layers [14]. However, being limited to a single ANN model (CNNs) limits the perspective of these studies on how model-specific computations contribute to this correspondence.

### Contributions of Low-level Visual Features

The categorical organization of the ventral visual stream can be partially explained by the low-level image properties of different object categories [26]. Luminance and contrast alone suffice to categorize images into abstract concepts with significant accuracy [27]. Decoding of visual information from EEG/MEG signals also depends on visual regularities in the stimuli: low-level visual features dominate object decodability early after stimulus onset (50 ms), with more abstract semantic representations appearing at later times (>150 ms) [4]. However, it remains to be addressed whether brain-ANN temporal correspondence is driven by their shared transformation of visual to semantic representation.

## 3 Methods

### 3.1 Datasets

We analyzed two versions of the THINGS-EEG dataset, consisting of EEG data recorded during visual perception (Fig. 1A). The THINGS-EEG 1 dataset includes EEG data from 50 participants (see A.2 for further details) presented with images spanning 1,854 concepts, each presented for 50 ms in a rapid serial visualization paradigm (RSVP) Fig. 1B) [28]. The THINGS-EEG 2 dataset includes 10 participants presented with images spanning 1,654 concepts, each presented for 100 ms with 100 ms of interstimuli interval [29]. From both datasets, we considered a subset of 200 unique images, each representing a distinctive concept.

**Figure 1.**
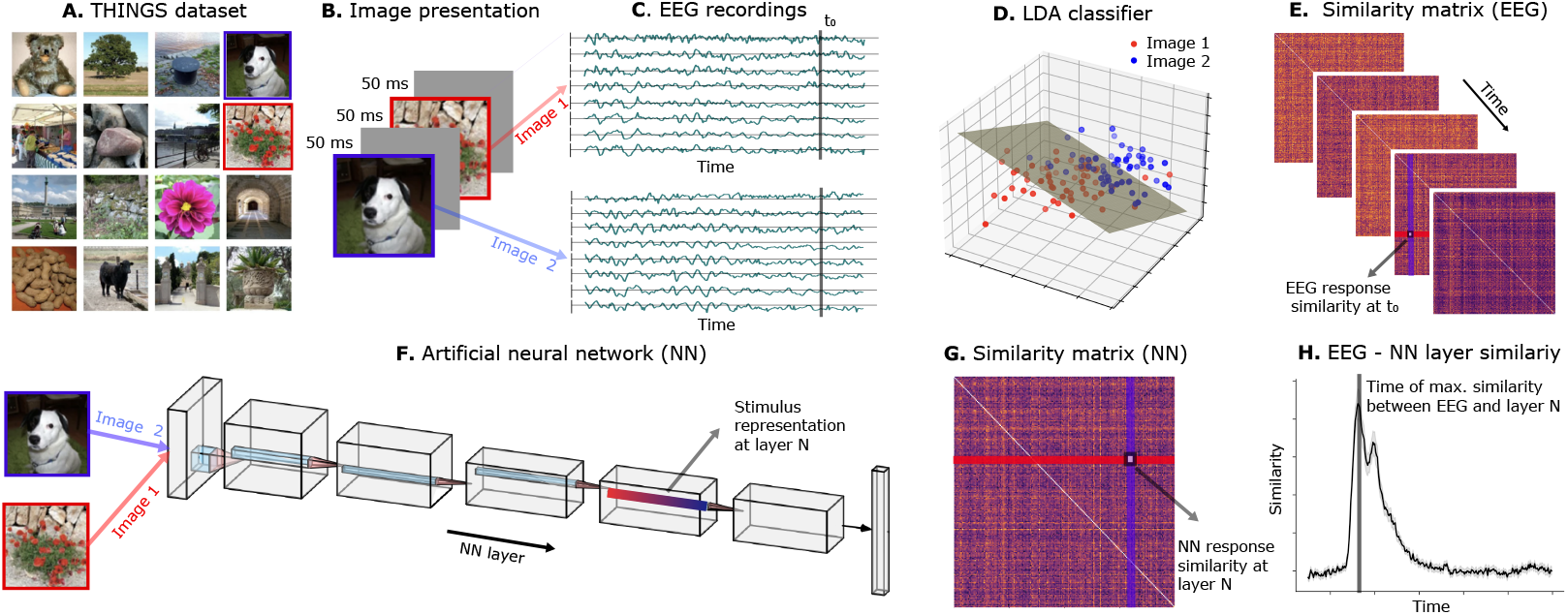
Overview of analysis methodology. A) Public domain images similar to the ones from the THINGS dataset (from PublicDomainPictures.net). B) Images were presented for 50 ms and 100 ms in THINGS-EEG 1 and THINGS-EEG 2, respectively, with inter-stimulus intervals of the same duration. C) For each image, EEG signals were acquired at 64 channels at different scalp locations. D) An LDA model was trained to classify all pairs of images at each time point using EEG signals as features. E) EEG RDMs were obtained using LDA accuracy as a distance metric. F) Images were used as inputs to different ANNs and the activations at all layers were extracted. G) For each ANN model and layer, RDMs were constructed using the cosine distance between activations. H) For each time point, RDMs from EEG and each ANN model were compared using Spearman correlation as similarity metric.

### 3.2 Data Pre-processing

Pre-processing of the recorded EEG signals (Fig. 1C) included a Hamming windowed finite impulse response filter (0.1 Hz high-pass and 100 Hz low-pass), re-referencing to the average reference, and down-sampling to 250 Hz. EEG data were preprocessed by baseline correction, where the mean of the pre-stimulus interval was subtracted for each trial and channel separately. For all this operations, we used MNE-Python v.1.9.0 [30].

### 3.3 EEG Decoding Distance

We used regularized Linear Discriminant Analysis (LDA) implemented in *scikit-learn* [31] to classify object pairs based on EEG signals at each time point (Fig. 1D). For each image pair, trials were split using a 70/30 stratified split. A regularized Linear Discriminant Analysis (LDA) classifier was fit on the training set with tolerance 0.01 and evaluated on the held-out test set ^2^. The resulting accuracies were used to construct a 200 × 200 representational dissimilarity matrix (RDM) for each subject in each point in time (Fig. 1E).

### 3.4 Artificial Neural Network Architectures

Images corresponding to both datasets were fed to ANNs to obtain layer-specific activations (Fig. 1F) after reshaping to a common size of 224 × 224 pixels. ANNs included three models previously used by Gifford and colleagues: AlexNet (CNN with 5 convolutional layers followed by 3 fully-connected layers), MoCo (feedforward architecture with self-supervised training), and ResNet-50 (supervised feedforward neural network with 50 layers and shortcut connections between layers at different depths) [29]. All of these networks were pre-trained on object categorization using the ILSVRC-2012 training image partition. We also included VGG-19 and ViT. VGG-19 was pre-trained on the ImageNet dataset, which contains over 3 million images and their corresponding concepts. ViT was pre-trained on ImageNet-21k (14 million images, 21,843 classes) at resolution 224 × 224, and fine-tuned on ImageNet 2012 (1 million images, 1,000 classes) at resolution 224 × 224.

### 3.5 Artificial Neural Network Activations

The selection of layers for AlexNet, MoCo and ResNet-50 was based on Gifford et al [29]. For VGG-19, we extracted activations from the five pooling layers (*pool1* to *pool5*) and the second fully-connected layer (*fc2*). From ViT, we utilized the activations from all 12 layers, extracting the class token from each layer, which represents the entire input with 768 dimensions [32]. We then flattened each layer, then re-scaled it and finally performed an iterative poly kernel Principal Component Analysis (PCA) algorithm with 200 components. We computed the cosine distance between activations to create a 200 × 200 RDM for each layer (Fig. 1G).

### 3.6 Temporal Correspondence

We compared each subject’s RDM at a given time with the RDM of each layer across ANN architectures. Then, we investigated whether the peak correlation times correlated with the order of the layers in each architecture, and whether there were differences between architectures (Fig. 1H). All the correlations between architectures were calculated using Spearman rho and all the comparisons between architectures were done using Wilcoxon test. Finally, we calculated the slope of correlation between LDA-semantic correlation and LDA-activation correlation through layers or each architecture for each dataset. We employed the Mann-Whitney U to test if there were differences between both datasets in each layer.

### 3.7 Low-level Visual Features

We used the methods described by Lützow Holm et al. to quantify the statistics of low-level visual features [26]. We extracted the descriptors computed by Harrison [27] and added values for the red, green, and blue channels. Using a recursive procedure [26], we then determined the 8 most important variables: 3 related to spectral features (SFE), 2 related to object edges (OE), luminance, and the colors green and blue, followed by PCA with retention of the first principal component. By correlating the absolute difference between each pair of vectors, we obtained 200 × 200 RDMs for each dataset.

### 3.8 Semantic Distance

For the semantic distance between concepts, we used FastText embeddings ^3^ between concepts and computed the cosine distance between every pair of embeddings, resulting in 200 × 200 RDMs for each dataset [33]. This RDM was validated using another set of embeddings from a state-of-the-art large language model, NV-Embed-v2 [34] and Stella [35]. See A.1 for further details.

## 4 Results

### 4.1 Analysis of ANN representations

Figure 2A displays the correlation between RDMs corresponding to all layers for the THINGS-EEG 1 dataset, both within and between ANNs. The architectures exhibited a high internal correlation, except for the last layer of the ViT, which differed from the other layers within the same architecture and from the layers of the other architectures. There was also correspondence between the first four architectures in terms of the relative order of their layers. As a general trend, the correlation with image statistics decreased as a function of the layer (Fig. 2B), while the correlation with semantic information increased (Fig. 2C). ViT differed from the rest as the decline of its correlation with image statistics was subtler than the rest and the slope of the semantic correlation did not differ significantly from zero. Similar results are shown for THINGS-EEG 2 in Figs. 2D-F. Additional analyses using a held-out validation image dataset are provided in Appendix A.1, Figure A.1.

**Figure 2.**
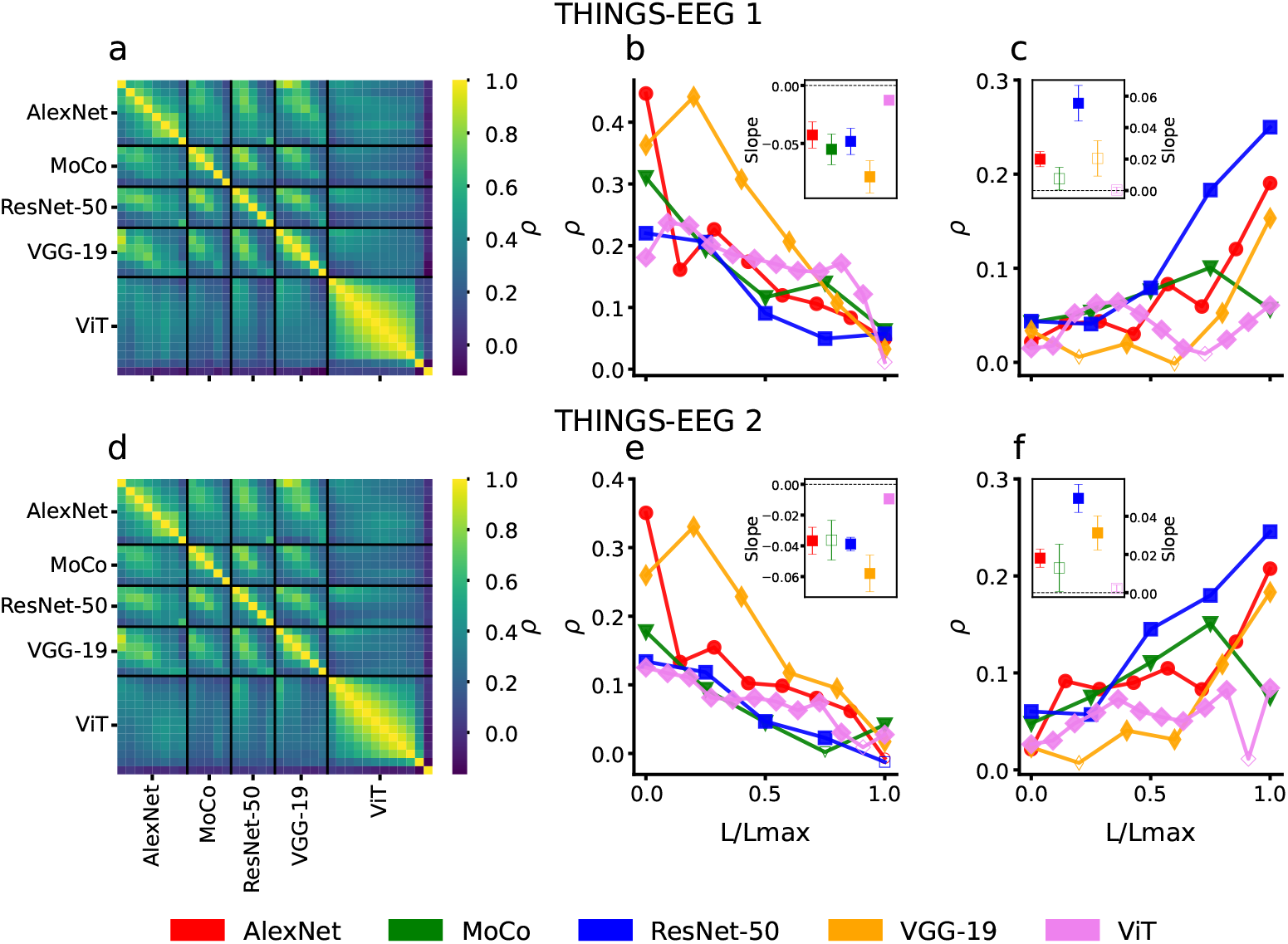
Transformation of visual to semantic representations in ANNs. A) Correlation between activations within and between ANNs for THINGS-EEG 1. B) Spearman correlation between RDMs based on low-level image statistics and ANN activations for THINGS-EEG 1. C) Spearman correlation between RDMs based on semantic similarity and ANN activations for THINGS-EEG 1. D, E, F) Same as panels A, B, C but for the THINGS-EEG 2 database. Full symbols indicate statistically significant Spearman correlation (p-value < 0.05).

### 4.2 Brain-ANN temporal correspondence

The time of maximum correlation (*T*_*max*_) between the ANN and EEG RDMs is shown in Fig. 3A (THINGS EEG 1) and B (THINGS EEG 2). *T*_*max*_ increased as a function of the layer, with exceptions for the last layer of VGG-19 and ViT in THINGS-EEG 1, as well as the last layer in both MoCo and ViT for THINGS-EEG 2. Also, the peak correlation between brain and ANN RDMs (*ρ*_*max*_) decreased as a function of the layer, except for the first layer of VGG-19 for both datasets and ViT in THINGS-EEG 1. Appendix Figures A.2–A.3 show the same *T*_max_ and *ρ*_max_ analyses computed over an independent validation set.

**Figure 3.**
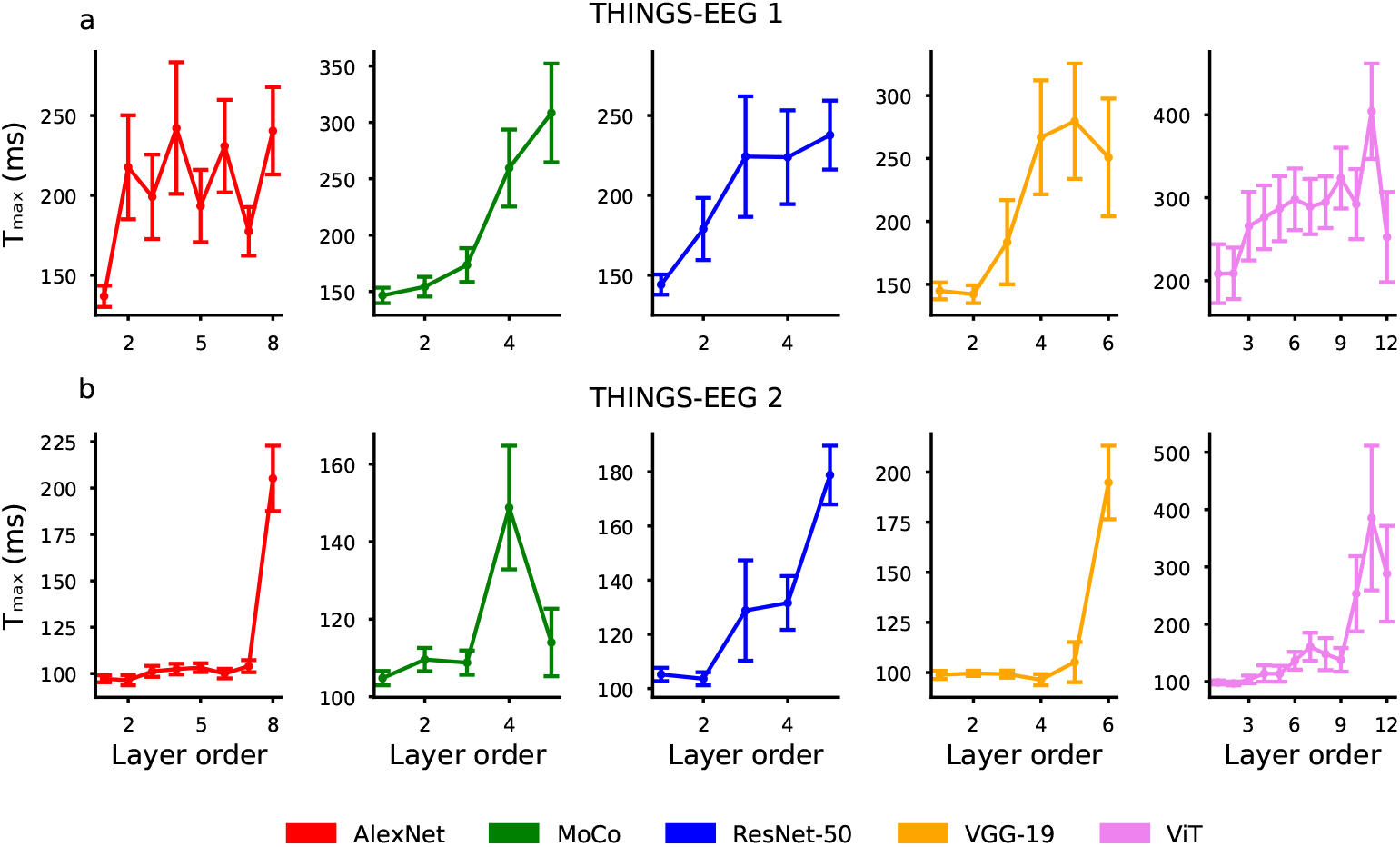
Time of maximum correlation between EEG and ANN activations. A) Time corresponding to the peak Spearman correlation between RDMs based on EEG and ANN activations (*T*_*max*_), for each layer of the models (layer order) and obtained using THINGS-EEG 1. B) Same as panel A, but for THINGS-EEG 2. Error bars correspond to standard error of the mean.

**Figure 4.**
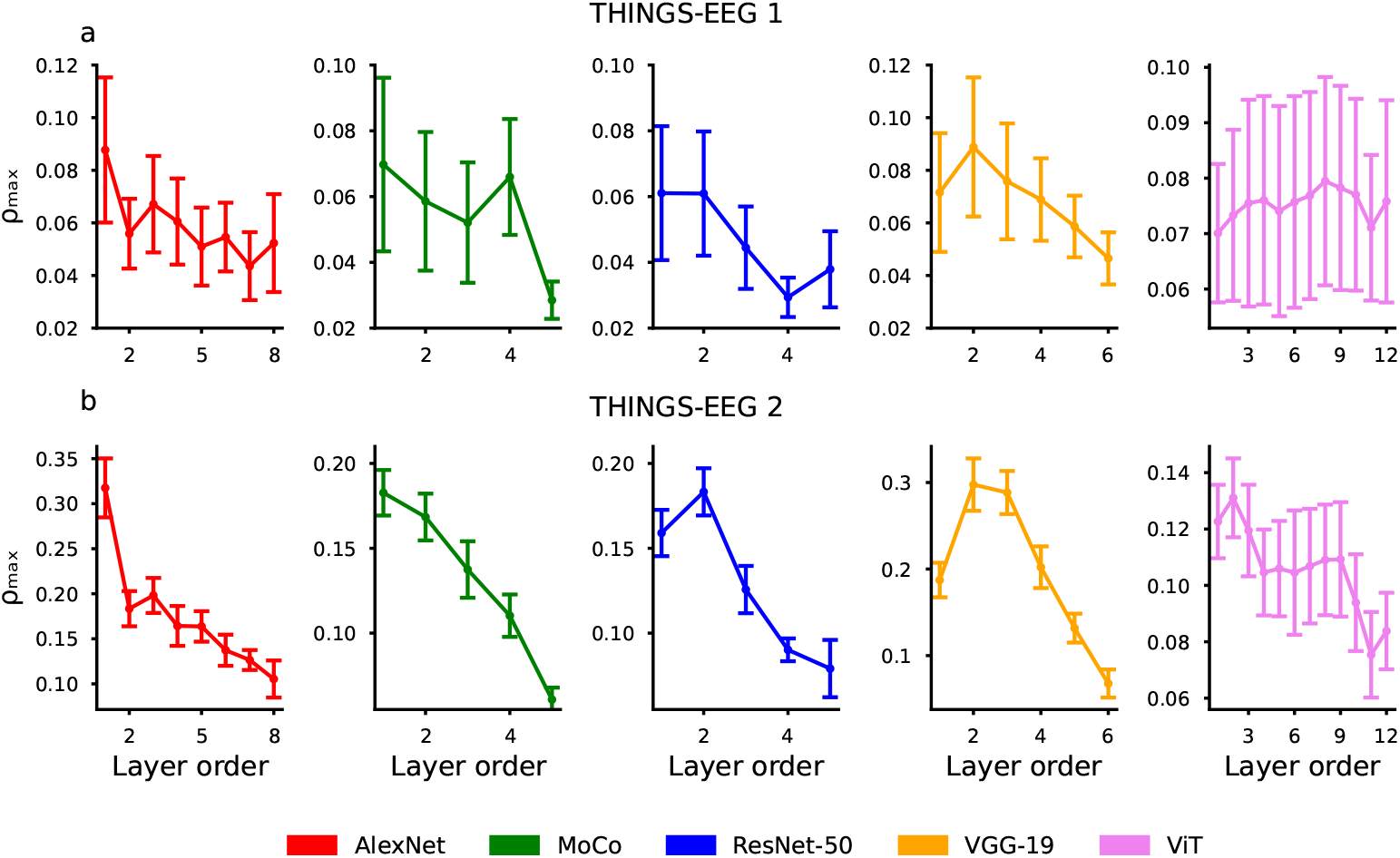
Peak correlation between EEG and activations. A) Peak value of the Spearman correlation between RDMs based on EEG and ANN activations (*ρ*_*max*_), for each layer of the models (layer order) and obtained using THINGS-EEG 1. B) Same as panel A, but for THINGS-EEG 2. Error bars correspond to standard error of the mean.

Next, we investigated the correlation between the layer order and the time of maximum correlation for each ANN (Figs. 5A and B for THINGS-EEG 1 and 2, respectively). For all models, there was a correspondence between *T*_*max*_ and layer order. Using the median value as a reference, ResNet-50 exhibited the highest time-layer correspondence, although there were no significant differences compared to the other architectures. Additionally, we investigated whether the peak correlation *ρ*_*max*_ behaved in a similar way (Figs. 5C and D). Except for ViT with THINGS-EEG 1, there was a negative correlation between *ρ*_*max*_ and layer order for all ANNs, indicating that the strength of brain-ANN correspondence decreased as a function of the layer. We found that *ρ*_*max*_ was closer to -1 and presented less variance for THINGS-EEG 2 compared to THINGS-EEG 1.

**Figure 5.**
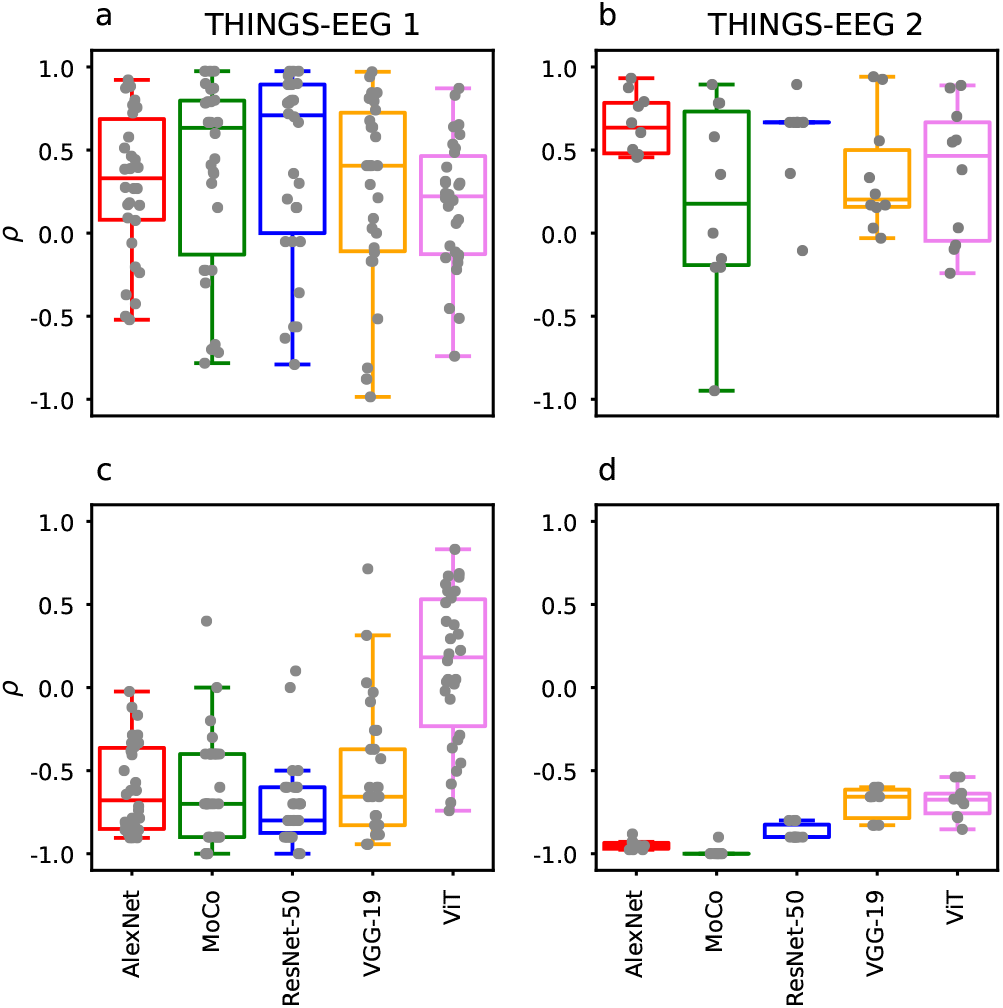
Monotonous behavior of *T*_*max*_ and *ρ*_*max*_ vs. layer order. A) Spearman correlation between layer order and *T*_*max*_ for THINGS-EEG 1. B) Same as in panel A, but for THINGS-EEG 2 dataset. C) Correlation between *ρ*_*max*_ and layer order for THINGS-EEG 1. D) Same as in panel C, but for THINGS-EEG 2.

### 4.3 Contributions of visual and semantic information

We then compared the RDMs obtained from image statistics and semantics with those computed from the EEG data (Fig. 6). We observed two significant peaks for the image statistics correlation in THINGS-EEG 1 (100 and 200 ms) and a very conspicuous peak for THINGS-EEG 2 (near 100 ms) followed by two smaller peaks at 200 and 300 ms. For THINGS-EEG 1, there was no significant correlation between the EEG and semantic RDMs. For THINGS-EEG 2, significant correlations between EEG and semantic RDMs were observed after the peak of correlation with the image statistics RDMs. As expected, the LDA curve for THINGS-EEG 1 was aligned with the visual component of the EEG, suggesting that for shorter stimuli, low-level features drove almost exclusively the discrimination between concepts. In contrast, the LDA curve for THINGS-EEG 2 had peak aligned with the visual component, but with a slower decay, as the semantic component was more relevant to distinguish image pairs. We also evaluated the correlation between the RDMs based on image statistics and semantic embeddings, without obtaining significant results (THINGS-EEG 1: r= -0.009, p-value < 0.05; THINGS-EEG 2: r = 0.011, p-value > 0.05). See Appendix Figure A.4 for similar results obtained in an independent validation dataset.

**Figure 6.**
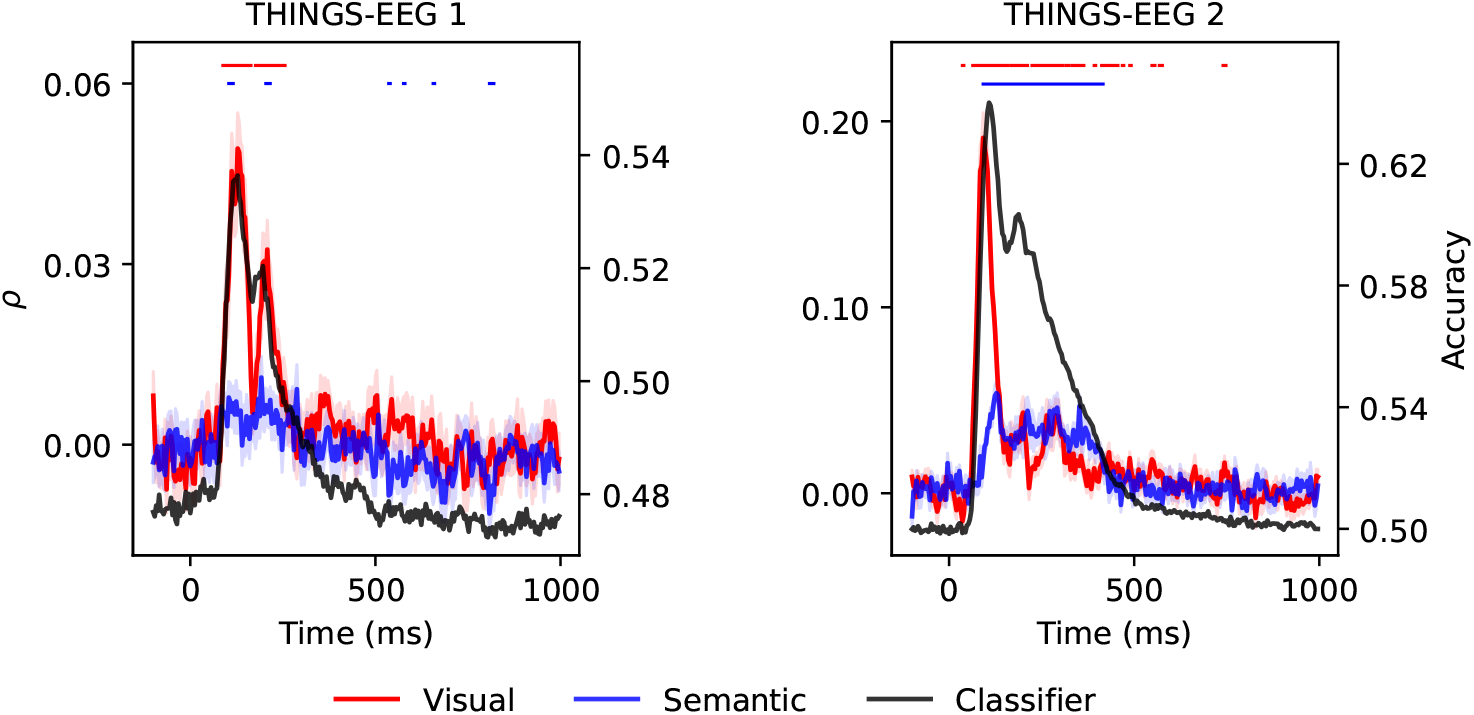
Contribution of low-level visual features and semantics to the EEG RDMs. Correlation between image statistics (red) and semantic (blue) RDMs, and the EEG RDM vs. time, for THINGS-EEG 1 (left) and THINGS-EEG 2 (right). The black curves correspond to LDA classifier accuracy. Lines above the plots indicate points in time with statistically significant Spearman correlation (red for statistics and blue for semantic, respectively; one sample t-test, p-value < 0.05). Shaded areas indicate standard error of the mean.

Finally, we computed the linear regression between EEG-semantic and EEG-activation correlations, and calculated the slopes for each layer of each architecture in THINGS-EEG 1 and 2 (Fig. 7). For AlexNet, Moco, ResNet-50 and VGG-19, THINGS-EEG 2 showed a significantly higher slope than THINGS-EEG 1 for some layers (one tailed t-test, p-value < 0.05). In these cases, brain-ANN correspondence was predicted by the semantic contribution to EEG responses across participants.

**Figure 7.**
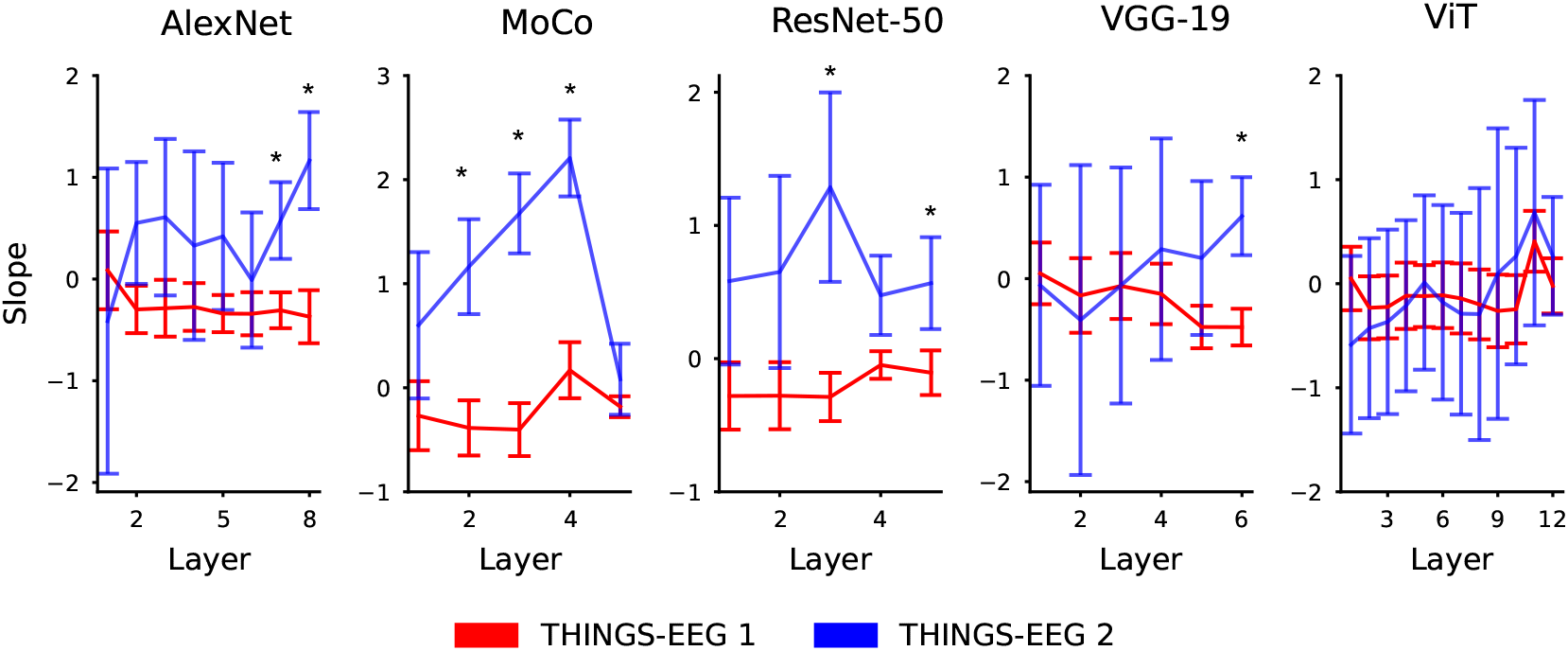
Regression of LDA-semantic vs. LDA-activation correlations. Slope of the best linear regression vs. layer order for all ANNs, comparing THINGS-EEG 1 (red) vs. THINGS-EEG 1 (blue). Asterisks indicate statistically significant differences between the slopes corresponding to both datasets (one tailed t-test, p-value < 0.05). Error bars correspond to standard error of the mean.

## 5 Discussion and Conclusions

We demonstrated a robust and consistent temporal correspondence between the temporal hierarchy of artificial neural network activations and human EEG responses during visual perception. Across multiple ANN architectures, we observed that early EEG components aligned with layers encoding low-level visual features, whereas later EEG components corresponded with deeper layers that capture semantic content. This alignment was modulated by stimulus duration, with longer exposures revealing stronger correspondence to semantic representations. Finally, for longer presentations times we observed a link between brain-ANN correspondence and the semantic contribution to EEG decoding (with the exception of the vision transformer). This result suggests that brain-ANN is driven by low-level visual representations for shorter stimulus duration, while semantic representations drive brain-ANN for longer durations, especially at deeper layers.

Our findings suggest that the observed brain–ANN reflects a shared computational principle: the progressive transformation from visual input to abstract semantic representations over time. Notably, even architectures without clear biological plausibility (e.g., ViT) exhibited this correspondence, supporting the idea that temporal alignment is driven more by representational hierarchy than by biological fidelity. This result is in agreement with fMRI and electrophysiological studies showing that architectural differences in AANs have little consequence in the prediction of brain responses during object recognition [16, 6].

Both the brain and ANNs process low-level visual information to obtain semantic representations. In the brain, the receptive field of neurons becomes increasingly larger and complex along the ventral pathway [36, 2]. Similarly, the hierarchical organization of CNNs results in units capable of invariant activation to specific objects [17]. Our analysis showed that this principle extends to other architectures: as layer order increases, activation similarity becomes more unrelated to low-level visual similarity and is increasingly predicted by semantic similarity. Since a similar progression is manifest in EEG responses, it is plausible that brain-ANN correspondence is primarily driven by the shared hierarchical transformation of visual to semantic representations.

We observed that the similarity between RDMs computed from EEG and AAN activations (*ρ*_*max*_) decreased as a function of the layer order. As shown in Fig. 6 and consistent with previous reports, the low-level visual features present a larger contribution to early EEG responses compared to semantic content [4]. Thus, this gradual reduction of *ρ*_*max*_ may parallel the shared shift from visual to semantic representations in the brain and ANNs. Moreover, Fig. 5 shows that *ρ*_*max*_ decreased at a faster rate in the dataset with shortest presentation times. The lack of significant semantic contribution to EEG responses observed in this dataset suggests that, in this case, brain-ANN correspondence could have been solely dominated by low-level visual features during the first ~200 ms after stimulus presentation (corresponding to the first layers of all models except ViT, as shown in Fig. 3. This result is aligned with known limits to post-perceptual processing for very short stimulus presentation times in the ∼`200 ms post-stimuli [37] (see also Fig. 6).

While the ViT model behaved similarly to other ANNs in terms of its temporal correspondence with EEG activity, it also presented some distinct features. As seen in the off-diagonal elements of Figs. 2A and D, a hierarchical correspondence existed between the intermediate layers of all ANNs, with the exception of the ViT. This result is in agreement with previous reports and is indicative of transformer-specific representations [38], which reinforces the independence of brain-ANN correspondence with regards to the idiosyncratic computational features of individual network architectures. The ViT model also behaved differently (i.e. non-monotonously) regarding the relative contribution of visual and semantic information, which could stem from the more uniform nature of representations across ViT layers [39].

In conclusion, our findings support the view that artificial neural networks resemble the brain not because they replicate specific low-level operations, but because they share representational goals—most notably, the transformation of visual input into semantic abstractions. This perspective de-emphasizes biologically motivated interpretations of CNNs and challenges the explanatory power of model-specific mechanisms such as convolutions, spatial locality, or receptive fields. Future work should investigate the role of hierarchical organization in the emergence of brain-like representations by systematically comparing models with varying degrees of hierarchical structure, including shallow or non-hierarchical architectures.

## 6 Limitations

### EEG spatial resolution

The limited spatial resolution of EEG restricted our ability to localize the neural sources of observed effects onto specific anatomical substrates, limiting our analysis to the temporal aspects of brain-ANN correspondence. Future work using source-localized MEG or intracranial recordings could provide more fine-grained spatial insights.

### Variance in the training set and tasks

Generalization from THINGS to naturalistic or dynamic visual environments (e.g., videos, scenes) remains untested. In particular, the limited sub-set of images may constrain the generalizability of the observed temporal correspondences. Also, while we employed a range of ANN architectures, all were trained on variants of ImageNet and optimized for object classification. Thus, our findings may not extend to models trained on other tasks.

### RSA

Similarity in representational space does not necessarily imply that the compared systems encode the same features or use similar representational formats. RSA is also sensitive to low-level confounds in the stimuli, which may inflate apparent correspondences driven by shared biases rather than meaningful alignment [40]. Finally, RSA offers only an indirect view of the temporal dynamics or transformations that occur within or across systems.

### Semantic similarity

Word embeddings provide a language-informed approximation of semantic structure based on distributional statistics of word co-occurrence, but they may not fully capture the distinctions most relevant to human visual perception. While restricting the analysis to images with a single dominant concept helps mitigate this concern, caution is warranted when extending this approach to more complex stimuli with ambiguous or multi-object content. Future work could address this limitation by incorporating more nuanced descriptions derived from large language models with multimodal capabilities [41].

## A Technical Appendices and Supplementary Material

### A.1 Correlation using embedding models

Correlation between fastText and Stella for THINGS-EEG 1: PearsonRResult(statistic = 0.47, p-value = 0.0) Correlation between fastText and NV-Embed-v2 for THINGS-EEG 1: PearsonRResult(statistic = 0.53, p-value = 0.0) Correlation between NV-Embed-v2 and Stella for THINGS-EEG 1: PearsonRResult(statistic = 0.55, p-value = 0.0)

Correlation between fastText and Stella for THINGS-EEG 2: PearsonRResult(statistic = 0.41, p-value = 0.0) Correlation between fastText and NV-Embed-v2 for THINGS-EEG 2: PearsonRResult(statistic = 0.55, p-value = 0.0) Correlation between NV-Embed-v2 and Stella for THINGS-EEG 2: PearsonRResult(statistic = 0.56, p-value = 0.0)

### A.2 Datasets

THINGS-EEG 1 ^4^ This study was approved by the University of Sydney Ethics Committee, and informed consent was obtained from all participants. Data are licensed under the Creative Commons Attribution 4.0 International License (CC BY 4.0). Of the 50 subjects, we included the 30 with the highest peak LDA values, as many of the remaining participants exhibited notably poor or noisy signals.

THINGS-EEG 2^5^ This study was approved by the Ethics Committee of Freie Universität Berlin (conducted in accordance with the Declaration of Helsinki). License: Creative Commons Attribution 4.0 International (CC BY 4.0) Repository: Open Science Framework

All validation tests were conducted using the dataset from Grootswagers et al. [42], which consists of EEG recordings from 16 adults viewing 200 visual objects presented at 20 Hz and then 5 Hz on a screen. The study was approved by the Ethics Committee of the University of Sydney. EEG recordings and images used as stimuli are available in the Open Science Framework repository ^6^. Repository: Open Science Framework. Image License: Creative Commons Attribution-NonCommercial 4.0 International (CC BY-NC 4.0)

### A.3 Implementation details

All computations were performed using Python 3.11.7 on a Ubuntu 22.04 Intel® Core™ i9-10900 CPU @ 2.80GHz × 20, Mesa Intel® UHD Graphics 630, 64 GB RAM with 1 TB storage. Our code is publicly available at https://anonymous.4open.science/r/0BF6/README.md.

**Figure A.1:**
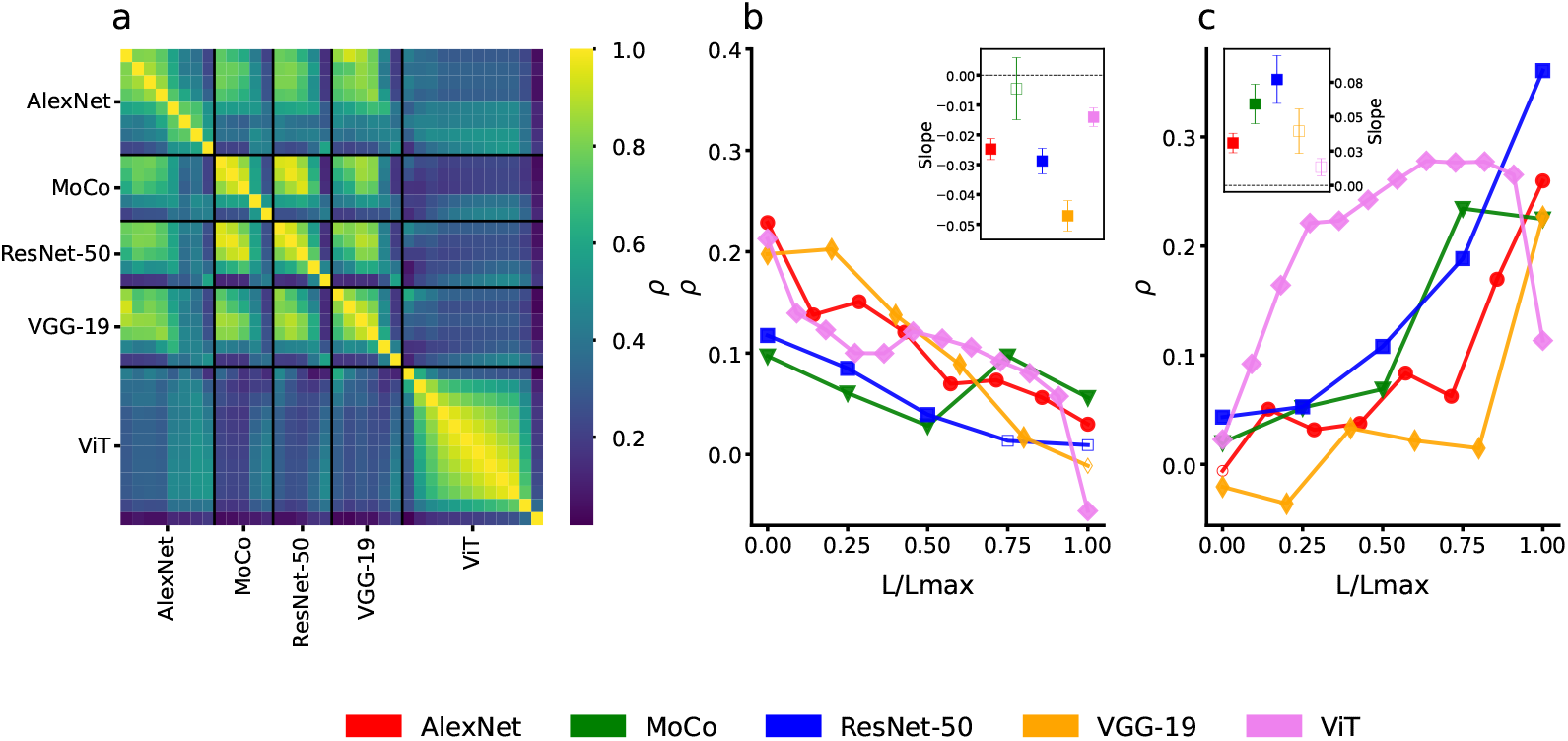
Comparison of activations for each layer of the architectures. A) Heatmap of correlation between RDMs for each layer of each architecture using the images of the validation dataset. B) Spearman correlation between image statistics RDM and activations RDM for the validation dataset. C) Spearman correlation between semantic correlation RDM and activations RDM for the validation dataset. Insets: slope for each curve with standard deviation. Full markers indicate statistical significance from zero (p-value < 0.05).

**Figure A.2:**
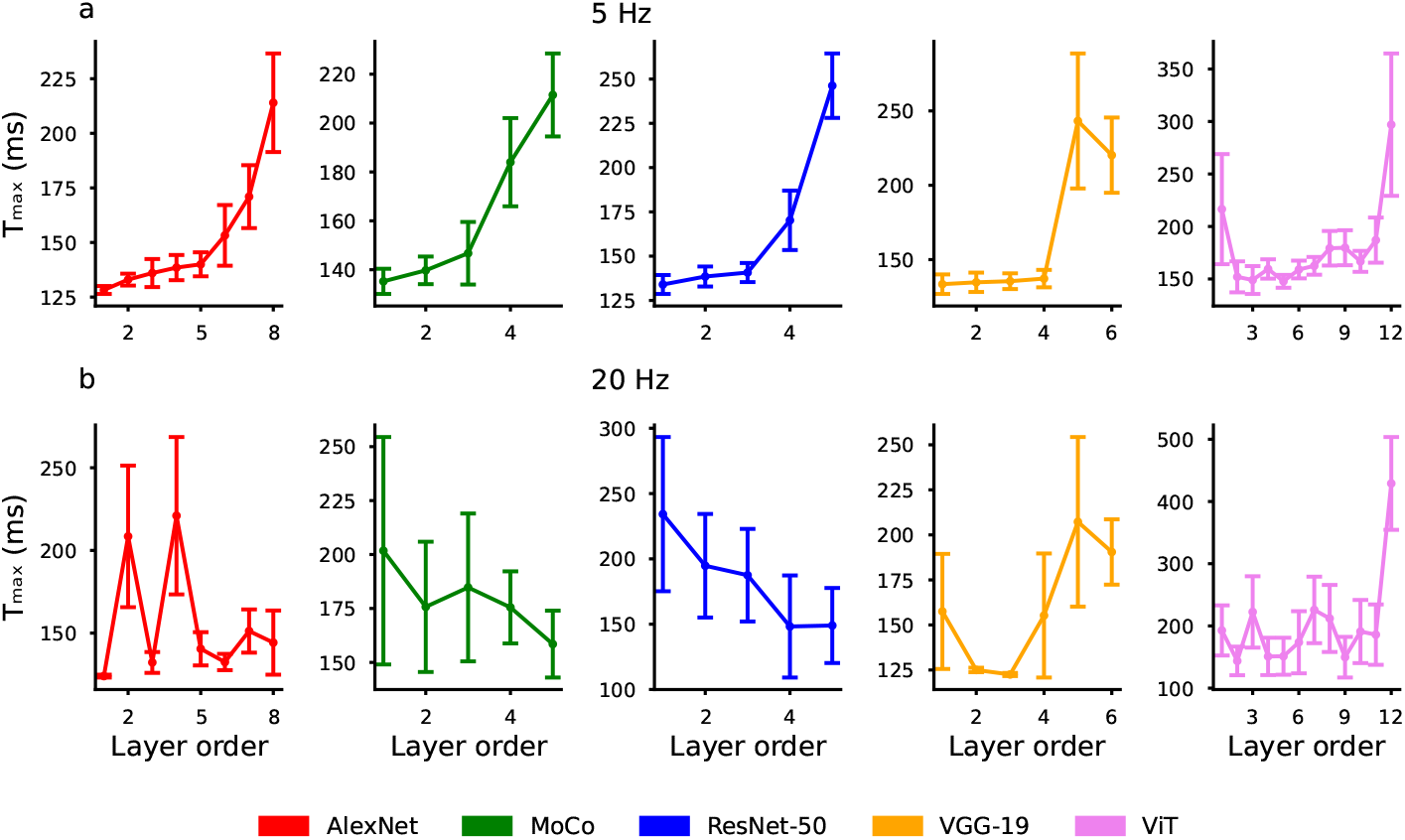
Time of maximum correlation between EEG and activations. A) Average time of maximum Spearman correlation between RDM of each layer and LDA for the validation dataset of 5 Hz. B) Same as A, but for the validation dataset of 20 Hz. Error bars correspond to standard error of the mean.

**Figure A.3:**
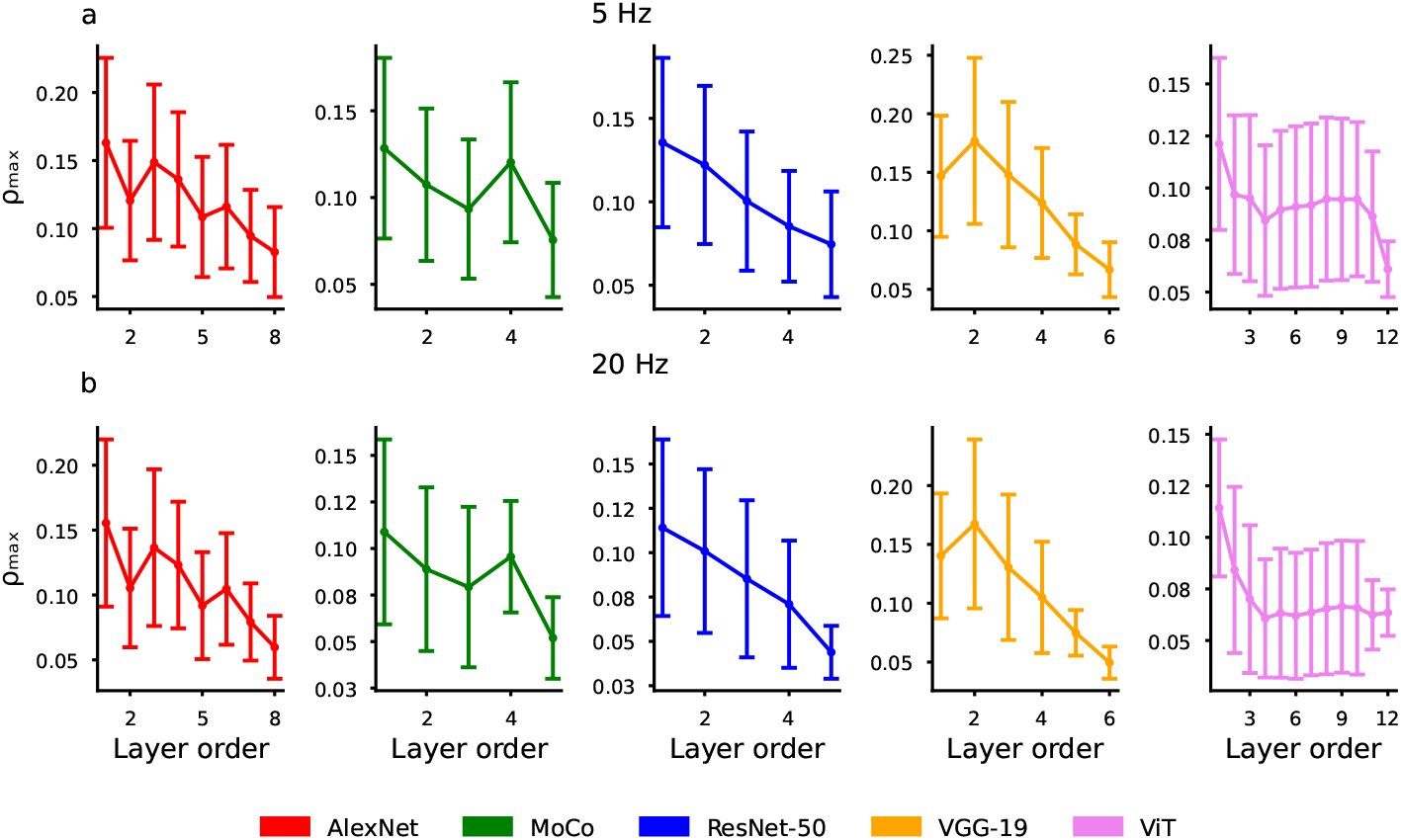
Value of maximum correlation between EEG and activations. A) Average maximum value of Spearman correlation between RDM of each layer and LDA for the validation dataset of 5 Hz. B) Same as A, but for the validation dataset of 20 Hz. Error bars correspond to standard error of the mean.

**Figure A.4:**
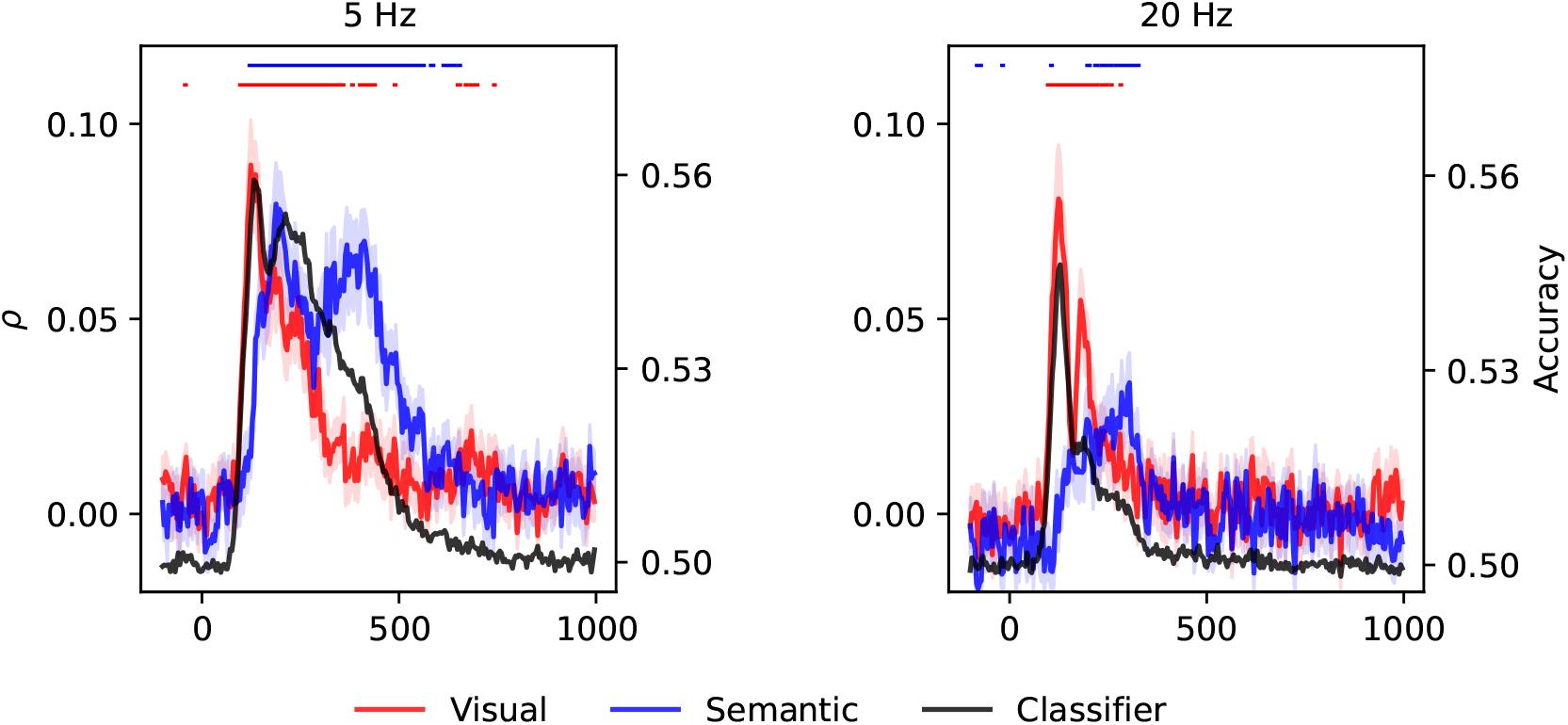
Contribution of low-level visual features and semantics to the EEG RDMs. Correlation between image statistics (red) and semantic (blue) RDMs, and the EEG RDM vs. time, for the 5 Hz (left) and 20 Hz (right) validation dataset. The black curves correspond to the LDA classifier used in each dataset. Lines above the plots indicate points in time with statistically significant Spearman correlation (red for statistics and blue for semantic, respectively; one sample t-test, p-value < 0.05). Shaded areas indicate standard error of the mean.

### Footnotes

2 Refer to the code for further details A.3

3 Training Corpus: Common Crawl (600 billion tokens). 2 million word vectors. 300 dimensions embeddings

4 https://openneuro.org/datasets/ds003825/versions/1.2.0

5 https://osf.io/3jk45/

6 https://osf.io/a7knv/

